# What determines the FDC polarization and GC size in affinity maturation

**DOI:** 10.1101/2020.09.08.287144

**Authors:** Zishuo Yan, Hai Qi, Yueheng Lan

## Abstract

Germinal center (GC) is a particular biological structure produced for affinity maturation in the lymphoid follicle during the T-dependent immune response and is an important component of the humoral immune system. However, the impact of morphological features of the GC on antibody production is not clear. According to the latest biological experiments, we establish a spatiotemporal stochastic model to simulate the whole self-organization process of the GC including the appearance of two specific zones: the dark zone(DZ) and the light zone (LZ). We find that the development of light and dark zones in GC serves to maintain an effective competition among different cells and promote affinity maturation. On the other hand, by varying the GC size, a phase transition is discovered, which determines a critical GC volume for best performance in both the stochastic and the deterministic model. This critical volume is determined by the distance between the activated B Cell Receptor(BCR) and the target epitope of the antigen. The conclusion is confirmed in both the 2D and the 3D simulations and explains partly the variability in the GC size.

**Author summary:** Germinal center (GC) is an important component of the humoral immune system, which supports antibody affinity maturation and the generation of immunity memory. However, the impact of special morphological features of the GC on antibody production is not clear. According to the latest biological experiments, we establish a spatiotemporal stochastic model to simulate the whole self-organization process of the GC. We use the mixing index of different B cells to quantitatively describe the polarization in GC. With the increase of the mixing index, the affinity of plasma cells decreases gradually, even GC might collapse. Therefore, the development of light and dark zones in GC serves to maintain effective competition among different cells and promote affinity maturation. On the other hand, by varying the GC volume, a phase transition is discovered, which determines a critical GC volume for best performance in both the stochastic and the deterministic model. This critical volume is determined by the distance between the activated B Cell Receptor (BCR) and the target epitope of antigen. The conclusion is confirmed in both the 2D and the 3D simulations and explains partly the variability in the GC size.

## Introduction

It has been known for a long time that hosts face a virtually unlimited number of antigens, so the innate immunity alone is not enough for a full-scale disease prevention. Therefore, adaptive immunity is required in defending against various antigens [1]. Germinal center (GC), first described by Walther Flemming in 1884 [2], is a main structure to produce specific immunity, which comes into being in secondary lymphoid organ, e.g. spleen, lymph node, pharyngeal tonsil. According to a series of experiments, researchers found, after pathogens invade the human body, the affinity of antibodies increases dramatically over time. This phenomenon is known as affinity maturation [3], which takes place mainly in GC through a series of physiological events including clonal expansion, high rate mutation of immunoglobulin, positive selection and cyclic reentry [4–6, 8].

To describe the reactions occurring inside the GC, and the roles played by different cells, MacLennan [4] introduced a classic model in 1994, based on which experimental biologists further revealed detailed dynamics in GC in order to explain the rapid maturation of antibody affinity. At the same time, theorists also established a series of mathematical models [13–15, 19], which compared the simulation and the experimental results to justify or modify model hypothesis. For example, in 1993, the deterministic model built by Kepler and Perelson [12] concluded that during the immune response, it is necessary for centroblasts(CBs) to travel back to the DZ from LZ in order to accelerate affinity maturation. However, these simulations are all deterministic, which fail to capture the stochastic nature of the real immune response.

Up to now, many details of the immune response remain unclear, such as why GC has the structure of light and dark zones? How B cells interact with T cells and what factors determine whether B cells differentiate into plasma or memory cells? For reality, more and more theoretical biologists begin to use stochastic models to describe the immune response. For example, Meyer-Hermann incorporated two-photon imaging data into his simulation of cell movement and migration to check the role of chemokines in GC formation [18, 20, 22]. He also had built a stochastic model describing morphological properties of the GC reaction, revealing necessary conditions for the development of DZ in the first phase of monoclonal expansion [16]. Later, two mechanisms are proposed to explain the rapid affinity maturation: the first is the introduction of a refractory time to limit the interaction between B cells and Follicular Dendritic Cells (FDCs), and the second is a competition between B cells for the help from the Follicular Helper T Cell(Tfh). Both mechanisms help achieve good affinity maturation in silico, which is consistent with experiment [17]. Recently, he proposed how the B cells go out of the GC and broadly neutralize antibodies with extensive simulations [21, 24].

In biological experiments, it has long been observed that GCs have similar structures and roughly the same volume. But there are few experiments and simulations to analyze the underlying principle behind morphological features of the GC, such as the appearance of the light or the dark zone and the constancy and the variability of the GC size. these problems are obviously important, because the immune system has evolved over a long period of time to develop GC into the current form, certainly in order to utilize as efficiently as possible different resources to acquire resistance to external pathogens. Therefore, an in-depth quantitative study on the GC morphology not only helps us understand the accelerated affinity maturation, but also suggests ways in controlling the GC formation.

In this article, we first build a stochastic model based on which the impact of GC morphology on affinity maturation is quantitatively studied. One interesting question is why most groups of FDCs are polarized in GCs. From the simulation, we see that the CXCL12-expressing reticular cells (CRCs) in DZ and FDCs in LZ together separate somatic hypermutation(SHM) and mitosis from clonal selection, which serves to maitain the competition of specific B cells and thus bestow efficient affinity maturation. Next, we check the influence of the GC size on affinity maturation. Simulation results show that the GC has an optimal volume for affinity maturation, determined by a phase transition at a particular volume size beyond which the affinity of plasma cells is greatly improved. In addition, this phase transition is first order, which is clearly seen in a deterministic model derived from the stochastic one. More detailed study shows that the phase transition point is determined by the distance in the gene space between the activated B cell and the target epitope antigen in lymph nodes. The final basic question is whether a 2D model can be used to represent the real GC reactions, so we extend the model to 3D for comparison and find a positive answer.

## Materials and methods

### Two models of the GC dynamics

In this paper, we establish a stochastic model as an extension of the one proposed in an earlier article [16] being summarized in Appendix A. The simulation is carried out with an accelerated version of the Gillespie algorithm [32]. More details are considered in the current model together with some newly revealed regulatory biochemical reactions. For example, we consider the entry and leaving of the Tfh cells in the GC and check the dynamic equilibrium of the Tfh cells in the GC. Once fully activated B cells enter the primary follicle, we deploy 100 activated T cells in paracortex, which ensures that the T cells entering the GC accounts for 5% of the total number of cells in the GC. These activated T cells actively respond to the CXCL13 secreted by FDCs [37], the concentration of which is inversely proportional to the distance from the FDC. With this attraction, these T cells gather around the primary follicle but are prevented from entering by bystander B cells [29, 30, 38]. Thus, ICOSL expression by follicular bystander B cells, engagement with ICOS signaling optimizes phosphoinositide-3 kinase (PI3K)-dependent pseudopod dynamics, which promotes T cells persistent motility at the T-B border and enables T cells to cross the border and enter the primary follicle. According to a latest research from Professor Hai Qi, PD-1, another B7 family molecule, though inhibitive, is highly expressed by T cells. They found that PD-1 inhibits the entry of T cells into GC by suppressing PI3K, which was induced by the PD-L1 expressed on follicular bystander B cells [31]. Thus, in our model, we consider activities of these co-inhibitory molecular pairs, leading to a dynamic equilibrium of the T cells.

### The stochastic model

We use a rectangular grid to represent the entire GC, which has a spacing of 10µm (Fig.1A), providing a platform for different cells to migrate randomly. We assume that the same type of cells cannot occupy the same grid point, while different types of cells can. In the model, CBs and CCs are regarded as the same type. The state of a cell is described by a vector containing many elements, such as spatial location, protein concentration, etc. During the GC evolution, each possible state change is regarded as a reaction. When a reaction takes place, the corresponding vector element of the cell will be updated. The signal integrator protein (SIP), a hypothetical protein that can regulate the immune behavior of different B cells in the simulation (Appendix A). The parameters and initial conditions of the model are shown in Table 2.

**Table 1.** Symbol Table.

| symbol | meaning | value |
| --- | --- | --- |
| $n_0$ | The total number of FDC sites | 80 |
| $i$ | The level of affinity | $0 \sim 9$ |
| $CC_{iFDC}$ | The number of CCs with affinity level $i$ that are binding with FDCs | - |
| $n_{FDC}$ | FDC sites that have not been occupied by CCs | - |
| $T_0$ | The total number of activated Tfh cells | 20 |
| $T$ | The number of Tfh cells that are not contact with CCs | - |
| $B_i$ | The number of CBs with affinity level $i$ | - |
| $m$ | The probability of no mutation | 0.5 |
| $m_2$ | The probability of each mutation moves forward or move backwards | 0.25 |
| $P_i$ | The total concentration of SIP with affinity level $i$ | - |
| $TR_i$ | The number of CCs with affinity level $i$ that have been rescued by T cells | - |
| $FR_{iT}$ | The number of CCs with affinity level $i$ that are binding with Tfh cells | - |
| $R_c$ | The probability of CCs' recycling into the dark zone | 0.8 |
| $SUMB_i$ | The sum of the numbers of CCs and CBs with affinity level $i$ | - |
| $P_i/SUMB_i$ | The SIP concentration of each B cell at level $i$ | - |
| $R_1, R_2, R_3, R_4, R_5, R_6$ | Defined in Appendix A 2,3,4,5,6,7 | - |
| $CC_i$ | The number of CCs with affinity level $i$ | - |
| $EN_{CF}$ | The possibility of CCs encountering with FDC sites | 0.0001 |
| $k_r$ | The dissociation rate of CCs with FDCs | 0.01 |
| $k_a$ | The degradation rate of ingested antigens in CCs | 1 |
| $Ag_i$ | The ingested antigens by CCs with level of affinity $i$ | - |
| $FR_i$ | The number of CCs with affinity level $i$ that had contacted with FDC sites | - |
| $EN_{CT}$ | The possibility of CCs encountering with Tfh cells. | 0.0001 |
| $SP_i$ | The concentration of help signals in B cells with affinity level $i$. | - |
| $k_s$ | The degradation rate of help signals in $FR_i$ | 1 |
| $PB_i$ | The number of output cells of different affinity levels, that is, plasma cells and memory cells | - |
| $k_p$ | The degradation rate of total concentration of SIP with affinity level | 1 |

**Table 2.** Parameters of Stochastic Spatiotemporal Model.

| Parameter | 2D | 3D | illustration |
| --- | --- | --- | --- |
| Lattice spacing | 10$\mu\text{m}$ | 10$\mu\text{m}$ | Radius of B cell is about 5$\mu\text{m}$ |
| Radius of GC | 200$\mu\text{m}$ | 150$\mu\text{m}$ | |
| Number of seeder cell | 3 | 3 | About three cells enter the follicle. |
| Shape space dimension | 4 | 4 | Range of affinity (0~1) |
| Number of T cells | 100 | 2500 |  |
| T cell movement rate | 10.8$\mu\text{m}/\text{min}$ | 10.8$\mu\text{m}/\text{min}$ | - |
| Number of FDCs | 20 | 225 | - |
| Length of FDC arms | 20$\mu\text{m}$ | 20$\mu\text{m}$ | - |
| Basic rate of proliferation | 1/9h | 1/6h | - |
| CB movement rate | 1.5$\mu\text{m}/\text{min}$ | 1.5$\mu\text{m}/\text{min}$ | - |
| Basic rate of CB differentiation | 1/7h | 1/4.7h | - |
| CC movement rate | 5$\mu\text{m}/\text{min}$ | 5$\mu\text{m}/\text{min}$ | - |
| Basic rate of CC differentiation | 1/6h | 1/4h | - |
| Basic rate of FDC-CC dissociation | 1/0.1h | 1/0.1h | - |
| Basic rate of FDC-T dissociation | 1/0.5h | 1/0.5h | - |
| Synthesis Rate of SIP | 1h | 1h | After CBs obtain help signals from T cells |

**Fig 1.**
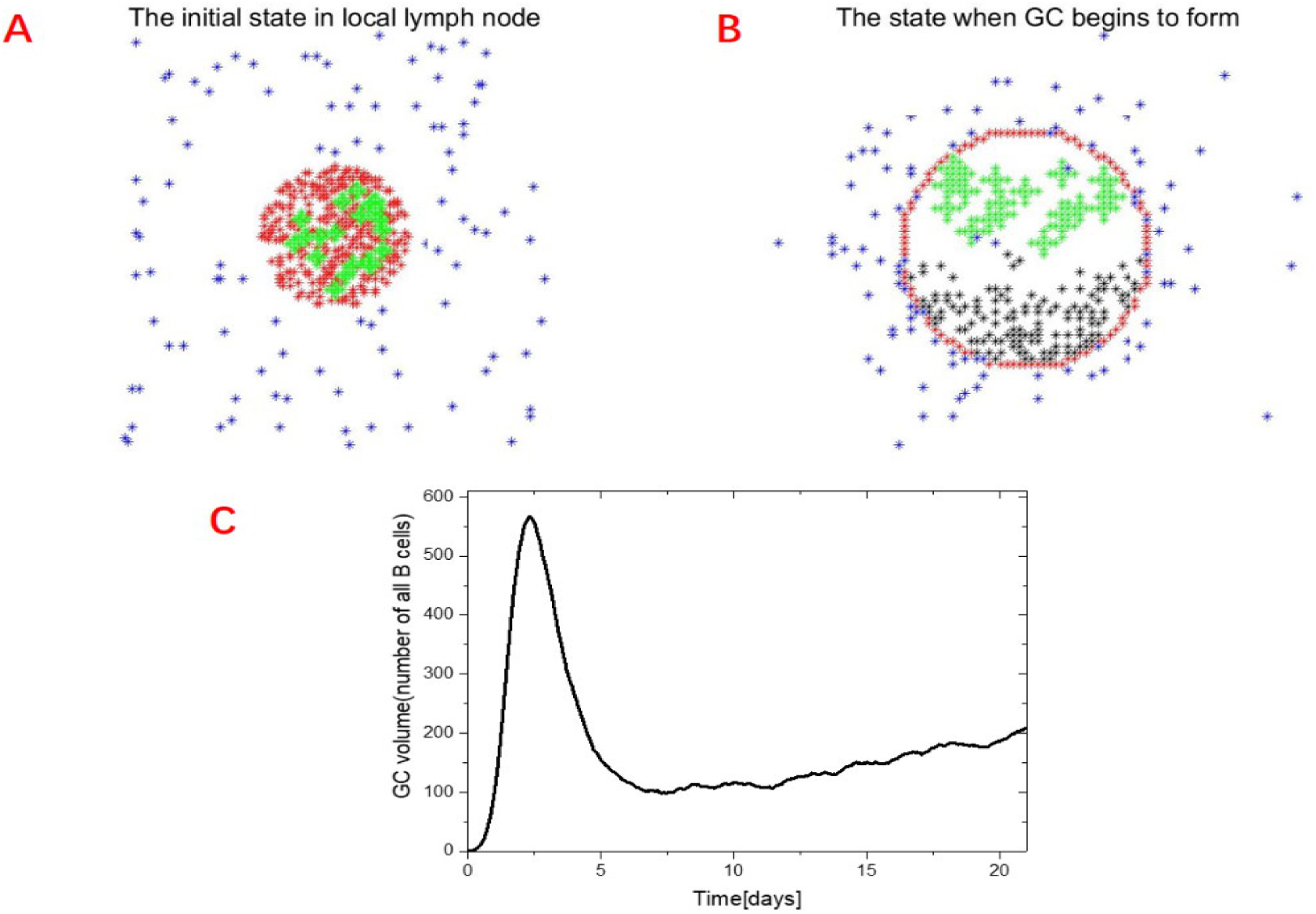
Cell configuration and GC volume in the simulation. (A) Screenshot of the simulated immune response at the initial time: naive B cells (red dots) fill the primary follicle, together with the FDCs (green crosses) in the center. The Tfh cells (blue dots) are wandering in the surrounding paracortex. (B) Tfh cells (blue dots) in the follicle move toward the follicle under the influence of chemokines. CBs (black dots) are rapidly dividing and form dark zone. FDCs are already in a polarized state in the LZ. Naive B cells that exist in primary follicles become boundaries as bystander cells (red dots). (C): Time course of the GC volume in the stochastic model.

### Cell motility

In our new model, the movement of B cells is dependent on chemokine concentrations. CCs could sense the concentration of CXCL13 secreted by FDCs. We assume that the concentration of CXCL13 is inversely proportional to the distance from the FDCs, 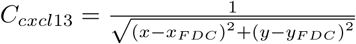 due to the proteolysis, where x,y is the location of CC, and *x*_*F DC*_, *y*_*F DC*_ is the average position of all FDCs. The attraction is realized by increasing the speed tending to the FDCs, which allows the cells leaving DZ for LZ to avoid apoptosis.

### T cells

When T cells are activated in the paracortex, the generation rate of ICOS and PD-1 is assumed to be 0.02/h and 0.01/h. When T cells are in contact with bystander B cells, due to the activation of ICOS and PD-1, the speed of T cells changes according to:

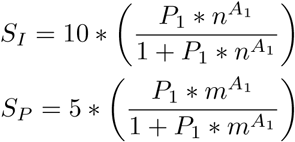

where *n* is the concentration of ICOS. *m* is the concentration of PD-1 in T cells. *P*_1_ = 4 × 10^*−*4^, *A*_1_ = 8. Due to the physical blockage of bystander B cells, the rate at which T cells enter a follicle is reduced. In our model, the speed of movement to the follicle is reduced by 5µm/min.

### Gene space

In the model, we use a four-dimensional lattice as the shape space to describe different gene types of antibodies and the target epitope of the antigen (Fig.2A). GC enriches the BCR phenotype by rapid division and proliferation of B cells with high frequency somatic mutations. In the stochastic model, the affinity of the antibody is assumed to double or reduce by half with each mutation of B cells. The epitope of the antigen is represented with a unique point – the origin *ϕ∗* =[0 0 0 0]. A mutation is represented by a jump to a nearest neighbor in the shape space. More explicitly, the affinity of an antibody can be defined as:

**Fig 2.**
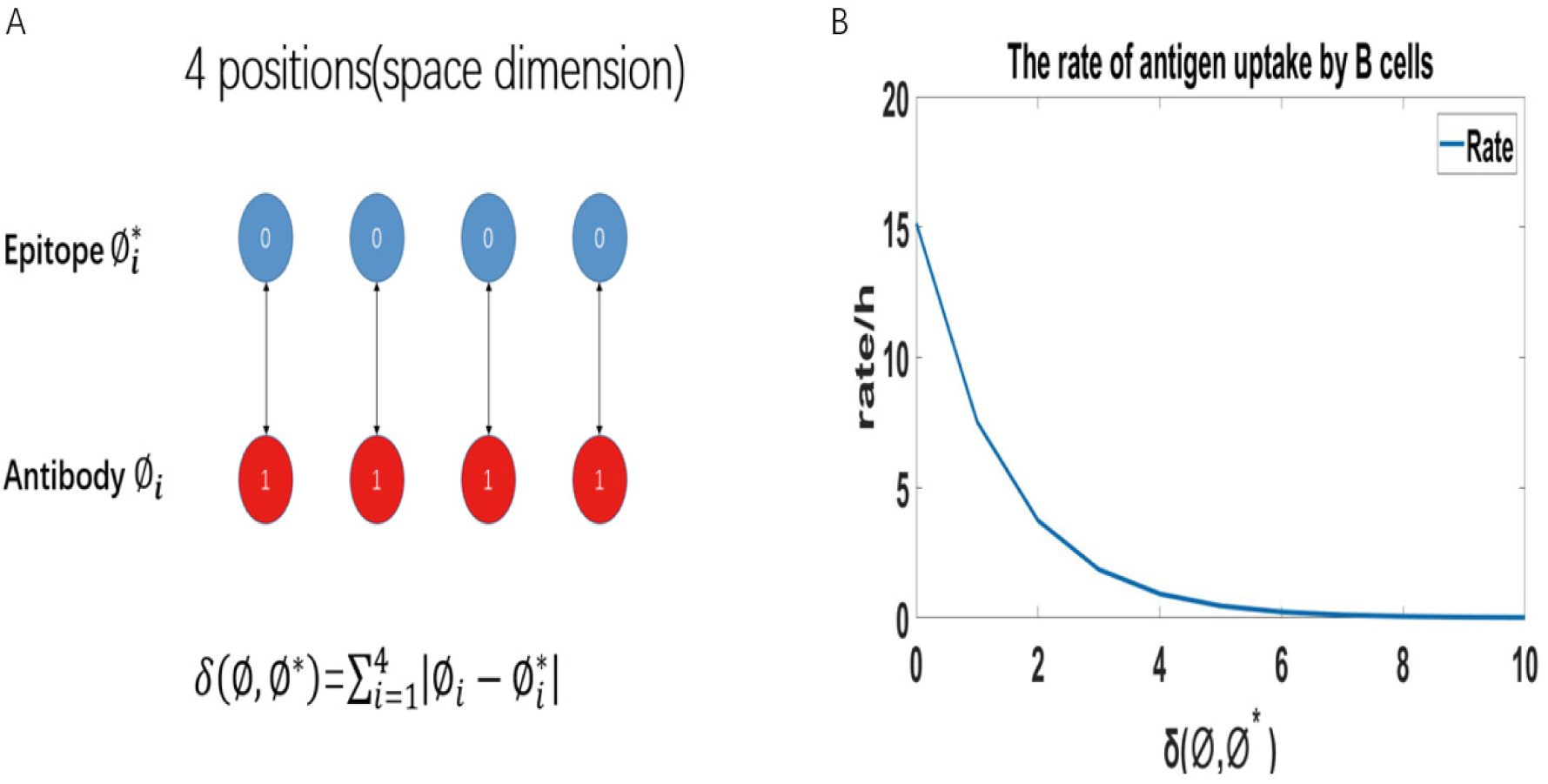
The distance *δ* in the gene space and how it changes the gene uptaking rate. (A) showing the relationship between rate of taking up Ag and *δ* (Appendix A7). (B) Antigens and antibodies are represented as points in the 4-dimensional lattice, and the define of *δ* (*ϕ, ϕ∗*).

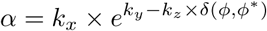

where *ϕ* is the coordinate of the BCR phenotype in the shape space, and the constant *k*_*x*_ = 0.0037, *k*_*y*_ = 5.6, *k*_*z*_ = 0.7. As can be seen from Fig.2B, the rate of taking up antigens from FDCs is higher when the affinity is higher. With the highest affinity, the CCs would take up one antigen in about every 4min.

### The deterministic model

Based on the stochastic model described above, a series of equations can be written down to describe the averaged dynamics in the GC. In the deterministic model, we set the affinity of BCR to 10 levels (0 *∼* 9) from low to high, which determines the binding rate of B cells to antigens. Correspondingly, the shape space is simplified a one-dimensional lattice with 10 sites (0 *∼* 9), over which cells with specific BCR phenotypes move randomly with equal probability in either direction except at the level 0 and 9 where moving in only one direction is possible. B cells rapidly clone at the early stage, producing a large number of offsprings with different affinities. In the deterministic model, we coarse-grain the states of B cells, assuming that B cells with the same affinity level have the same properties, such as the rate of division, the rate of differentiation, and the ability to bind to the antigen, etc. The initial B cells are assigned the zero affinity.

Next, we write down all the equations and explain each of them below.

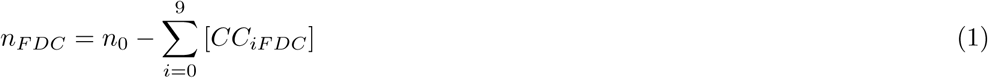

which describes the change of the FDC sites. *n*_0_(= 80) is the total number of FDC sites and *n*_*F DC*_ is the number of sites that are not been occupied by CCs. *CC*_*iF DC*_ is the number of CCs with affinity level *i* and binding with FDC.

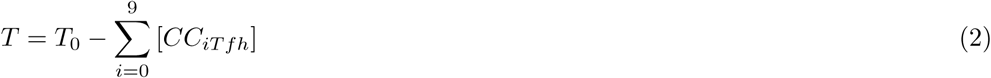

where *T*_0_(= 20) is the total number of activated Tfh cells and T is the number of Tfh cells that are not in contact with CCs. *CC*_*iT fh*_ is the number of CCs with an affinity level *i* and binding with Tfh cells.

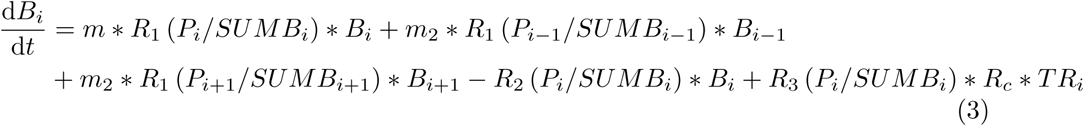

which describes the process of B cell proliferation and differentiation. *B*_*i*_ is the number of CBs with affinity level *i*. The probability of each mutation is 1 *− m*. The probability of each mutation moves forward or move backwards is *m*_2_. *P*_*i*_ is the sum of the concentration of SIP. *TR*_*i*_ is the number of CCs rescued by Tfh cells with an affinity level of *i*. These CCs either differentiate into plasma cells or recycle into the DZ to continue dividing. The *R*_*c*_(= 0.8) is the probability of CCs’ recycling into the DZ. *SUMB*_*i*_ is the sum of the numbers of CCs and CBs with an affinity level *i*, so the SIP concentration in each B cell at level *i* is *P*_*i*_*/SUMB*_*i*_, which then give the rates *R*_1_, *R*_2_, *R*_3_ defined in Appendix A2,A3,A4.

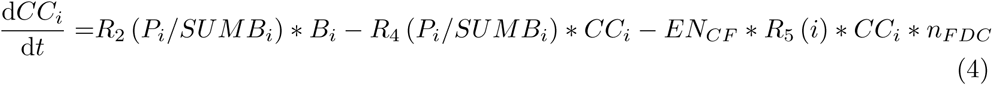

which describes the death of CCs and its interaction with FDCs. *CC*_*i*_ is the number of CCs with affinity level *i. EN*_*CF*_ (=0.0001) is the possibility of CCs encountering with FDC sites. The rates *R*_4_, *R*_5_ are functions of the SIP concentration and defined Appendix A5,A6.

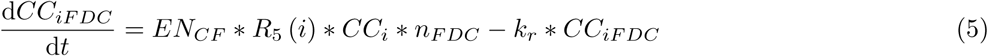

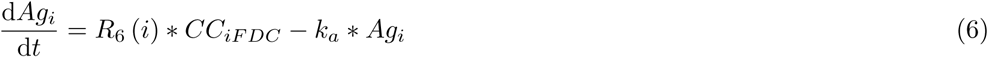

which describe the binding of CCs with FDCs and the change of the antigen concentration in *CC*_*i*_. *k*_*r*_(=0.01) is the dissociation rate of CCs with FDCs. *k*_*a*_(=1)is the degradation rate of ingested antigens. *Ag*_*i*_ is the concentration of the ingested antigens by CCs with affinity level *i. R*_6_ is a rate defined in Appendix A6.

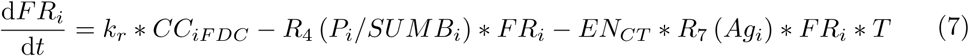

where *FR*_*i*_ is the number of CCs with affinity level *i* that had contacted with FDC sites and *EN*_*CT*_ (=0.0001) is the possibility of CCs encountering with Tfh cells.

In the deterministic model, the positive feedback of ICOS-ICOSL and CD40L–CD40 and the impact of PD-L1-PD-1 molecular pairs are not considered. Although these molecular pairs contribute to affinity maturation, the decisive factor is still the concentrations of antigens absorbed by CCs. Therefore, this model is not identical with the stochastic model and the impact of molecular pairs is not considered. Similar to what is in the stochastic model, the higher the antigen concentrations, the faster the binding rate of CCs with Tfh cells. The reaction rates *R*_7_ and *R*_8_ :

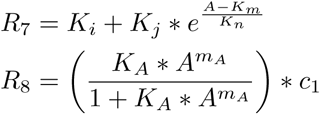

where A is the concentration of antigens, *K*_*i*_ = 10, *K*_*j*_ = 2, *K*_*m*_ = 10, *K*_*n*_ = 30. *m*_*A*_ = 8 is Hill’s coefficient, *K*_*A*_ = 6 × 10^*−*4^, *c*_1_ = 18.5.

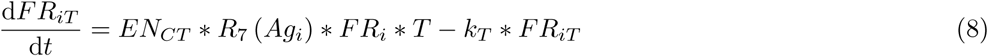

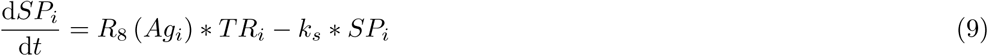

which describe the binding of CCs with Tfh cells and the help signals from Tfh cells. *k*_*T*_ (= 0.05) is the dissociation rate of CCs and Tfh cells. *FR*_*iT*_ is the number of CCs with affinity level *i* that are binding with Tfh cells. *SP*_*i*_ is the concentration of help signals in CCs. *k*_*s*_(= 1) is the degradation rate of help signals in *TR*_*i*_.

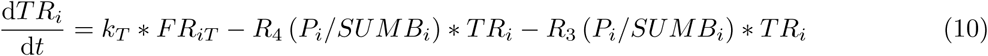

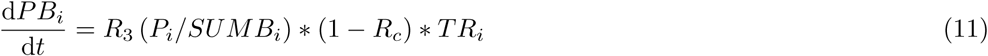

which describe the process of CCs differentiating into CBs and into memory or plasma cells. *PB*_*i*_ represents output cells with different affinity levels, namely plasma cells and memory cells.

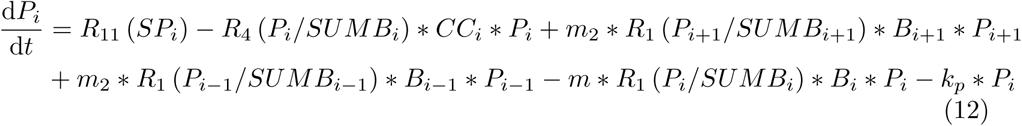

which describes the concentration changes of SIP. *k*_*p*_ is the degradation rate of total concentrations of SIP with affinity level *i*. In this model, the synthesis rate of SIP *R*_11_ is affected by the concentration of help signals in CCs:

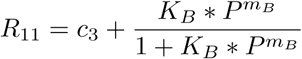

where P is the concentration of help signals and Hill’s coefficient *m*_*B*_ = 4. The constant *K*_*B*_ = 4 × 10^*−*4^, *c*_3_ = 1.

## Results

We use the spatiotemporal stochastic model based on the Gillespie algorithm to simulate the immune response in the GC within 21 days. When simulating in silico, we can easily change the morphology of GC and analyze its impact on affinity maturation [33, 34]. We simulate the GC space using a 120*120 lattice with the grid spacing being 10µm. The space was divided into two parts: the paracortex area and the primary follicle area (Fig.1A).

At the beginning of the simulation, activated T cells and B cells moved to the T-B boundary under the influence of chemokines. When these T cells meet an activated B cell, this B cell will be activated fully and move to the primary follicle to generate GC. At this time, FDCs will be polarized. Naïve B cells that exist in primary follicles make up their boundaries as bystander B cells. With the proliferation and differentiation of B cells (Fig.1B), LZ and DZ gradually appear in the GC, and the chemokines CXCL12 and CXCL13 play an important role in the whole process. The positioning of CBs in the DZ depends on the expression of CXCL12 receptors, CXCR4 [37]. When CBs differentiate into CCs, they express CXCR5, the receptor of CXCL13, which promotes CCs motility towards the LZ. (Fig.1A, Fig.1B: Simulation diagram in silico, T cells (blue), naïve B cells (red), FDC (green), CBs(black)).

### The FDC polarization

In the primary follicle, the FDC is in the central region. At the T-B border, there are many CXCL12-expressing reticular cells (CRCs) interlaced with FDCs. In the GC, FDCs stay in a polarized distribution, located in the LZ. In the DZ, there are mostly CRCs. This kind of distribution in the GC is observed across a range of species. Therefore, it is natural to ask, if polarization is eliminated, that is, if the light and dark zone disappear, and CBs and CCs mix with each other, what will happen to affinity maturation. In 2013, Bannard [39] used CXCR4-deficient GC B cells, confined in the LZ, to confirm that the differentiation from CBs to CCs is independent of their position. He also provided strong evidence that the spatial separation of LZ and DZ is critical to maintain an effective GC response. In biological experiments, CXCR4-deficient GC B cells have fewer accumulations of genetic mutations, mainly because there is less AID (activation-induced cytidine deaminase) in LZ than in DZ. The physiological function of AID is to introduce point mutation in the variable region during SHM. In our model, we may assume a uniform concentrations of AID in the GC to ensure an equal mutation rate everywhere, so that the impact of AID can be eliminated. The degree of mixing of CBs with CCs is described with a polarization index *I*_*p*_ that counts the number of CCs in a radius of 20µm around the CBs under investigation. By changing the distance between the FDCs and the GC center, the mixing index between CBs and CCs is varying. When FDCs are in the center of the GC, the mixing index is the greatest, and when FDCs are maximally polarized, CCs and CBs are separated to the maximum extent, which will form the LZ and DZ in GC.

As can be seen from the GC volume change in Fig.3A, on the tenth day, the number of B cells in the GC without partitions (Fig.3A Black line) is much lower than that with spatial polarization (Fig.3A Red line). It is because that B cells inside this type of GC do not compete properly. Because of the mixing, CBs may occupy the sites of the FDCs, causing excessively senseless competition between CBs and CCs. Due to the crowdedness in the lattice space, CCs cannot walk freely, which limits their chance of taking antigens from FDCs. As the mixing degree *I*_*p*_ is getting higher, the average affinity of the output plasma cells keeps decreasing (Fig.3B), which indicates the importance of the CRCs regulation in the DZ and polarization of FDCs in the LZ. Together, they separate SHM and mitotic process from selection, resulting in more reasonable competition between CCs.

**Fig 3.**
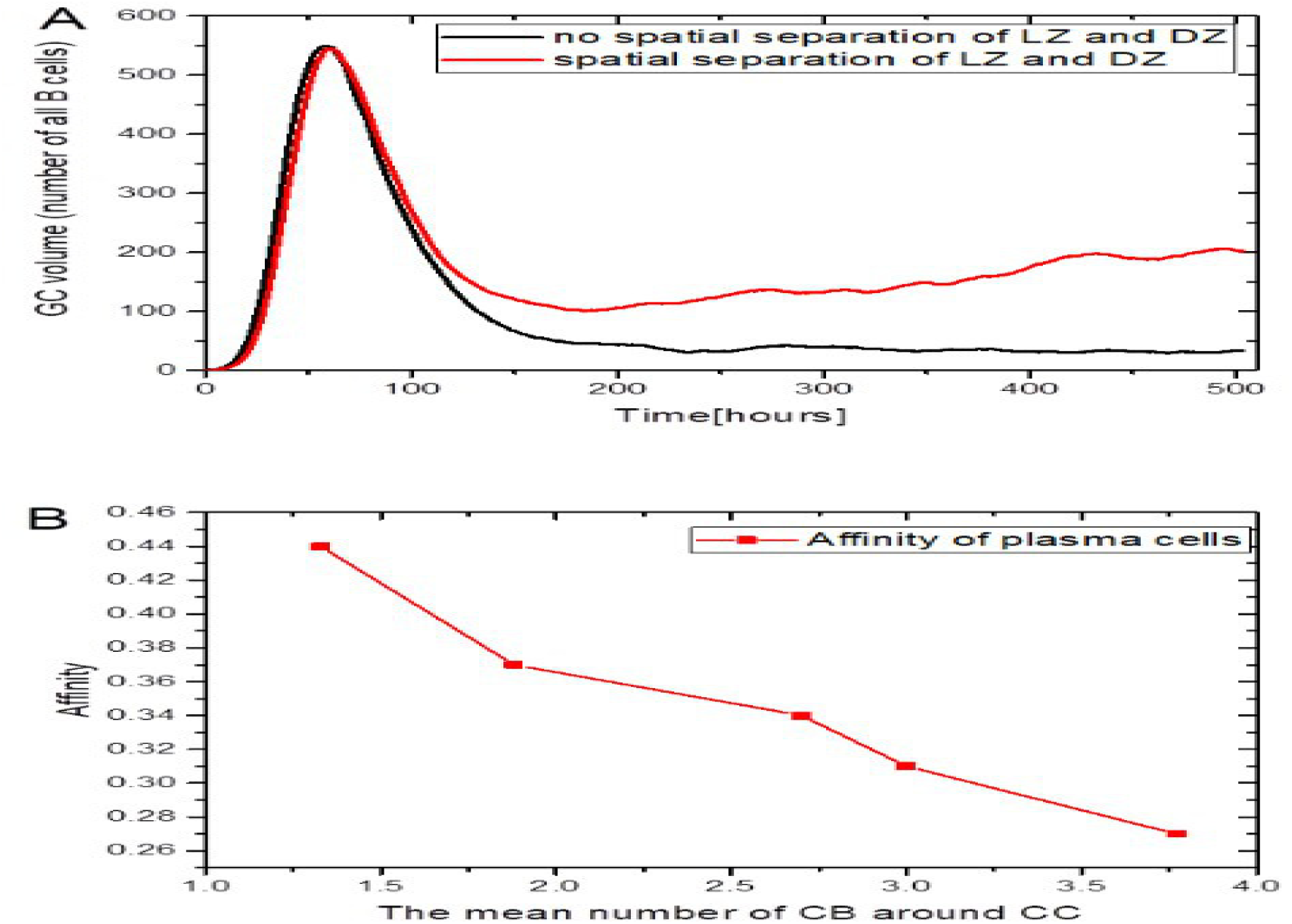
The influence of FDCs distribution in GC. A: Time course of the GC volume with spatial separation (Red, the mean number of CB around CC is 1.5), and without (Black, the mean number of CB around CC is 3.52). B: The affinity vs the polarization index *I*_*p*_

### Dependence of the affinity on the GC size

In the following, we discuss the impact of different GC volumes on affinity maturation. In the simulation, the density of FDCs and the number of T cells around GC remain constant. Six different radii, 50µm, 100µm, 150µm, 200µm, 300µm, 400µm, are used to carry out the computation which produce different affinities for the output cells. From Fig.4A, we can see from the simulation results that when the radius of GC is 200µm, it is the most suitable for affinity maturation, and there is a phase transition in affinity level when the radii is between 100µm and 150µm. When the phase transition occurs, the standard deviation of the affinity reaches the maximum, which indicates that the simulation results fluctuate greatly at this radius. With further increase of the radius, the affinity increases, while the standard deviation decreases, and the simulation results become more and more stable. When the radius exceeds the optimal radius, the affinity starts to decrease gradually with the increase of the radius. The reason for this decrease can be seen from Fig.4C. With the increase of the volume of GC, the accumulation of B cell mutation becomes more and more, a large portion of the mutation is farther and farther away from the target epitope of antigen, and the proportion of B cells with low affinity increases, so it becomes more difficult to select CCs with high affinity.

**Fig 4.**
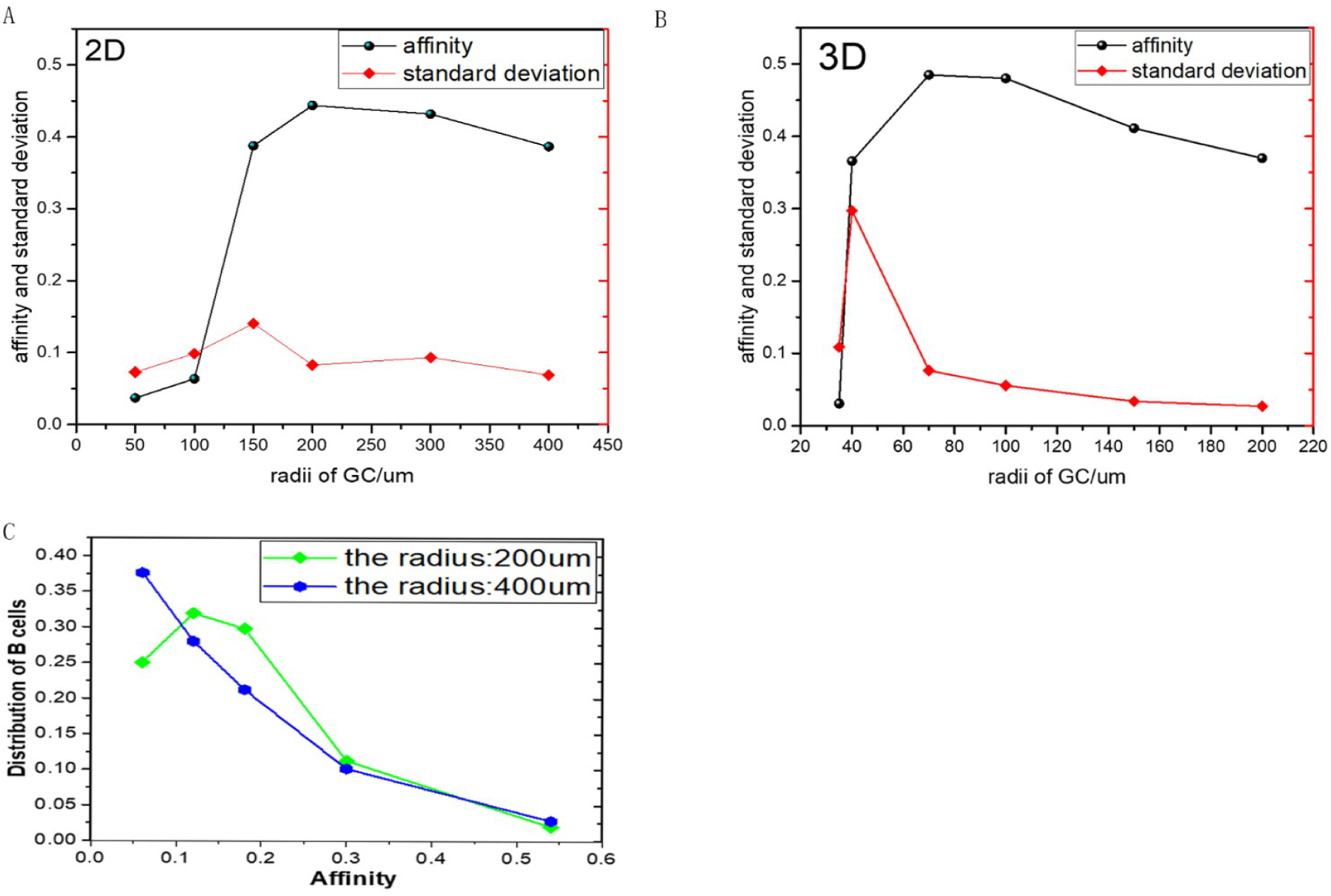
Average affinity and the standard deviation of output cells for different volumes of the GC in the 2D (A) and 3D (B) simulation. (C): Affinity distribution of B cells in the GC for two different radii on the third day, with a radius of 200µm (blue line) and 400µm (green line).

As shown in Fig.5, we simulate the development and affinity maturation of the GC with the deterministic equation. On the 21st day, the average affinity of all output cells was 0.37611. A similar phase transition can be seen in Fig.6C for the deterministic model, where the volume of the GC is controled by the concentration of SIP in the initial B cells. As a result, the initial SIP concentration could be used to represent the GC volume. The higher the concentrations of SIP, the larger the volume of the GC (Fig.6A). The jump of the affinity can also be observed when the concentration of SIP in naive B cells is 251, which clearly indicates a phase transition (Fig.6C). When the concentration of SIP is less than 251, the volume of GC will decay to zero with time, while when the concentration of SIP is higher than 251, GC will reach a stable volume. At 251, GC contained 110 B cells, which was the largest volume of GC (near the third day). Similar results are obtained when the critical radius is 100µm in the stochastic model (Fig.6D). The existence of the phase transition could also be explained qualitatively in a straightforward manner. If the number of CBs generated by division or proliferation is too small, there is not enough mutation accumulation to produce B cells with high affinity. As a result, these B cells cannot get enough help from Tfh cells and most of B cells will execute apoptosis, so the GC will eventually collapse. In order to achieve affinity maturation, the GC needs a sufficient number of B cells to undergo SHM. The phase transition point pins down the lower bound of this number in average.

**Fig 5.**
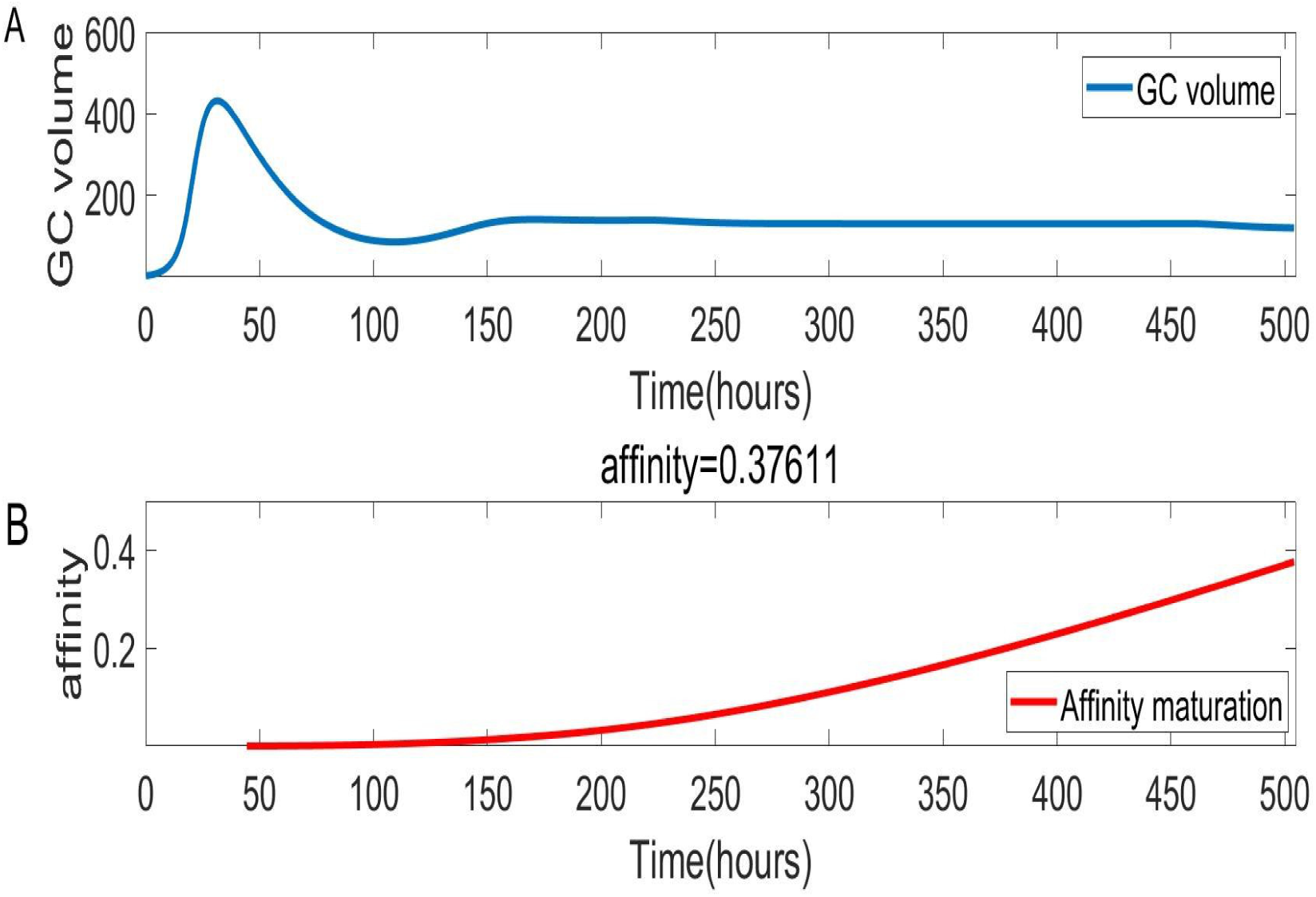
The evolution of the GC volume and the affinity. (A) the change in the volume of the GC over time, (B) the maturation course of the output cell affinity.

**Fig 6.**
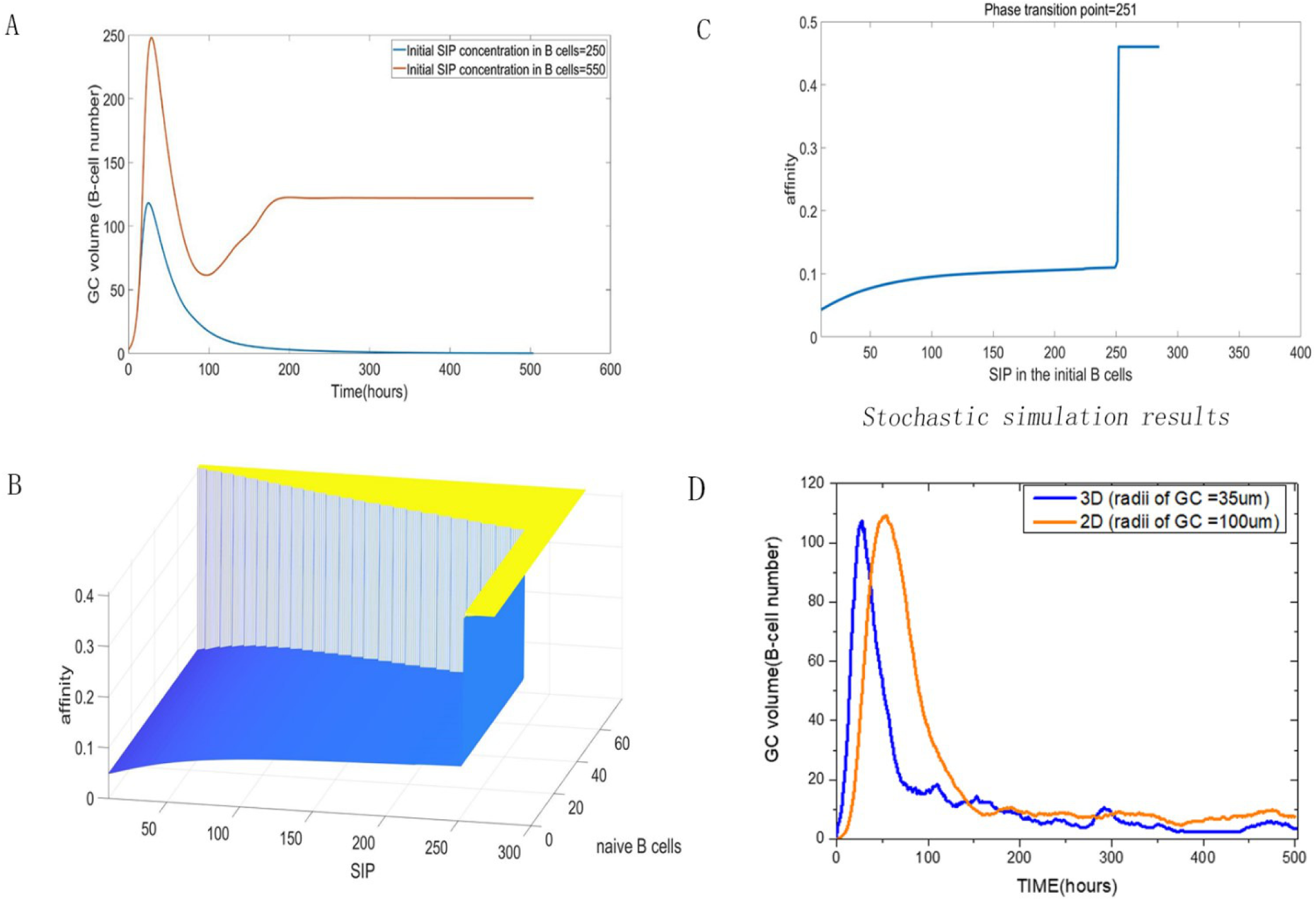
The phase transition observed in deterministic model. (A) The change of the GC volume with time for SIP=250 and 550 in the initial B cells. (B) A three-dimensional plot of the dependence of the average affinity of the output cells on the concentration of SIP in the initial B cells and the number of newly added B cells with higher affinity at the initial time. (C) The average affinity of the output cells is dependent on the concentration of SIP in the initial B cells. (D) Time course of the GC volume in the stochastic model.

Below, we will investigate what controls the critical number of B cells for the phase transition. In the stochastic model, the initial B cells are assumed to have a genotype [1 1 1 1], the distance between BCR phenotype and the target epitope of the antigen is *δ* (*ϕ, ϕ∗*) = 4. This distance plays an important role in determining the critical GC volume as will be seen next. If we set the initial BCR genotype to [1 1 0 0] and [2 2 2 2] respectively in the stochastic model, the distance to the target epitope is *δ* (*ϕ, ϕ∗*) = 2 and *δ* (*ϕ, ϕ∗*) = 8. As can be seen from Fig.7A and B, the affinities in these two cases do not reach the same level. When *δ* (*ϕ, ϕ∗*) = 2, even if the volume of the GC is very small, the average affinity of output cells is greater than 0.5, but when *δ* (*ϕ, ϕ∗*) = 8, the maximum affinity is 0.2, achieved at a radius as large as 400µm. Of course, by increasing the volume of the GC, it is possible to promote cell affinity. With *δ* (*ϕ, ϕ∗*) = 2 increasing to *δ* (*ϕ, ϕ∗*) = 4, the GC radius with the highest average affinity increases from 150µm to 200µm. Therefore, we may draw the conclusion that the critical GC volume depends on the distance *δ*(*ϕ, ϕ∗*), which may be checked experimentally.

**Fig 7.**
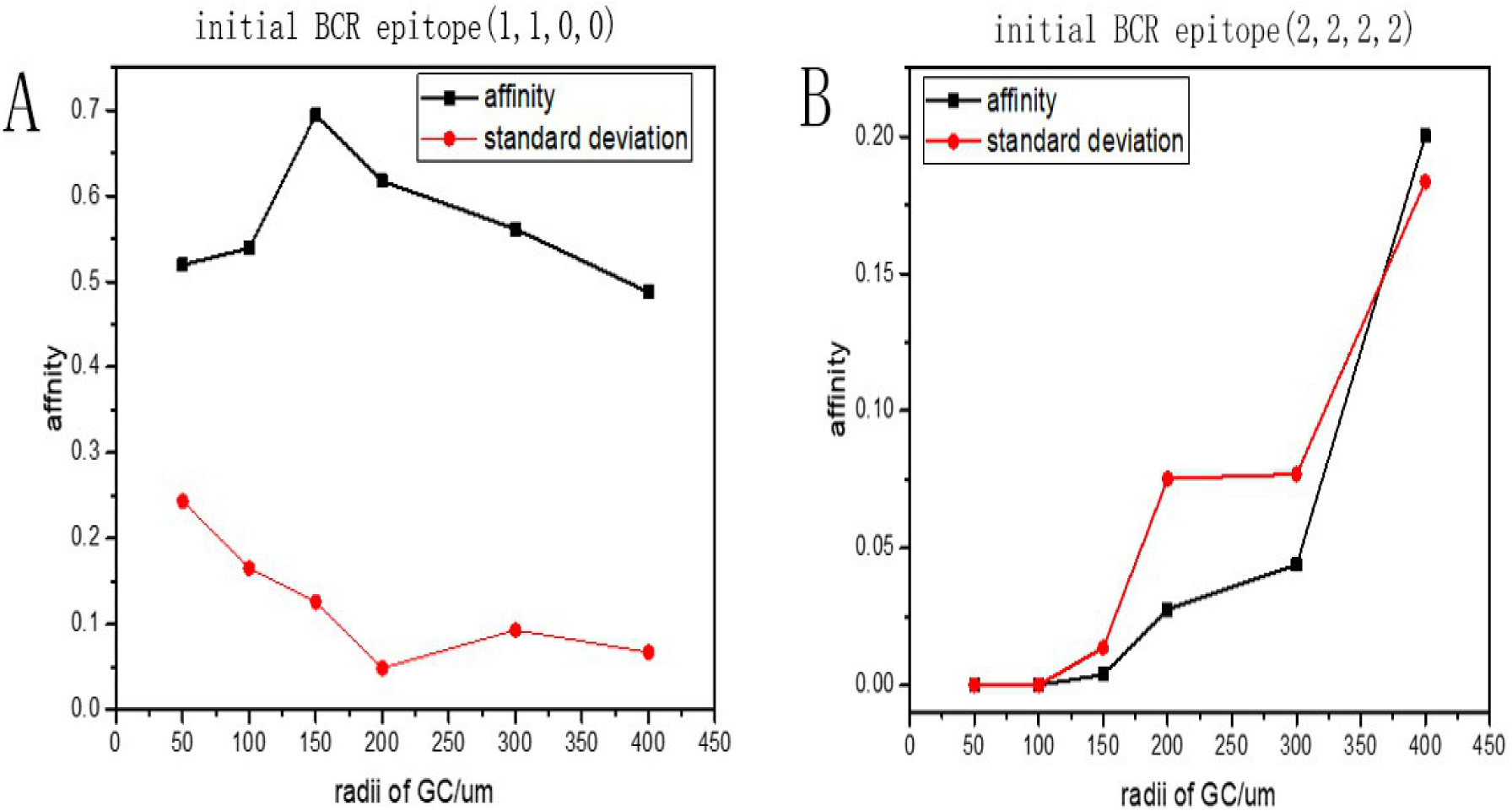
When the epitope is [0 0 0 0], the average affinity of output cells for different volumes. (A) The phenotype of BCR is (1,1,0,0). (B) The phenotype of BCR is [2 2 2 2].

When the volume of the GC is very small, it is important to find a mechanism to ensure that plasma cells with high affinity could eventually be produced. In biological experiments, it has been observed that GCs with different sizes grow in lymph nodes, between which cells migrate now and then. Therefore, It is reasonable to speculate that some B cells with high affinity may be able to move to smaller GCs to help them grow and promote the production of high affinity plasma cells. To verify this hypothesis, in both the deterministic and the stochastic model, we initially supply additional B cells with high affinity to help affinity maturation and check the affinity variation of the output cells. As can be seen from Fig.6B, in the deterministic simulation, there are two major factors that modulate the affinity level of the output cells: the volume of GC and the number of B cells with different affinity added at the initial time. Similar results could be obtained in the stochastic model. In one trial, ten B cells with an affinity of 0.24 are put to the follicle in addition to the original three B cells with an affinity of 0.06. From Fig.8A, we can see that the affinity of the output plasma cells is 0.35, much higher than 0.2, the affinity achieved without this additional input. However, if we change the affinity of the ten B cells from 0.24 to 0.49 (Fig.8B), the affinity of plasma cells is improved further to 0.5 and the standard deviation becomes much smaller. From the results of Fig.6B and Fig.8, it is reasonable to say that B cells with higher affinity that exit from GCs and enter other smaller GCs may help produce high affinity output cells, and thus contribute to the collective maturation in the whole lymphoid.

**Fig 8.**
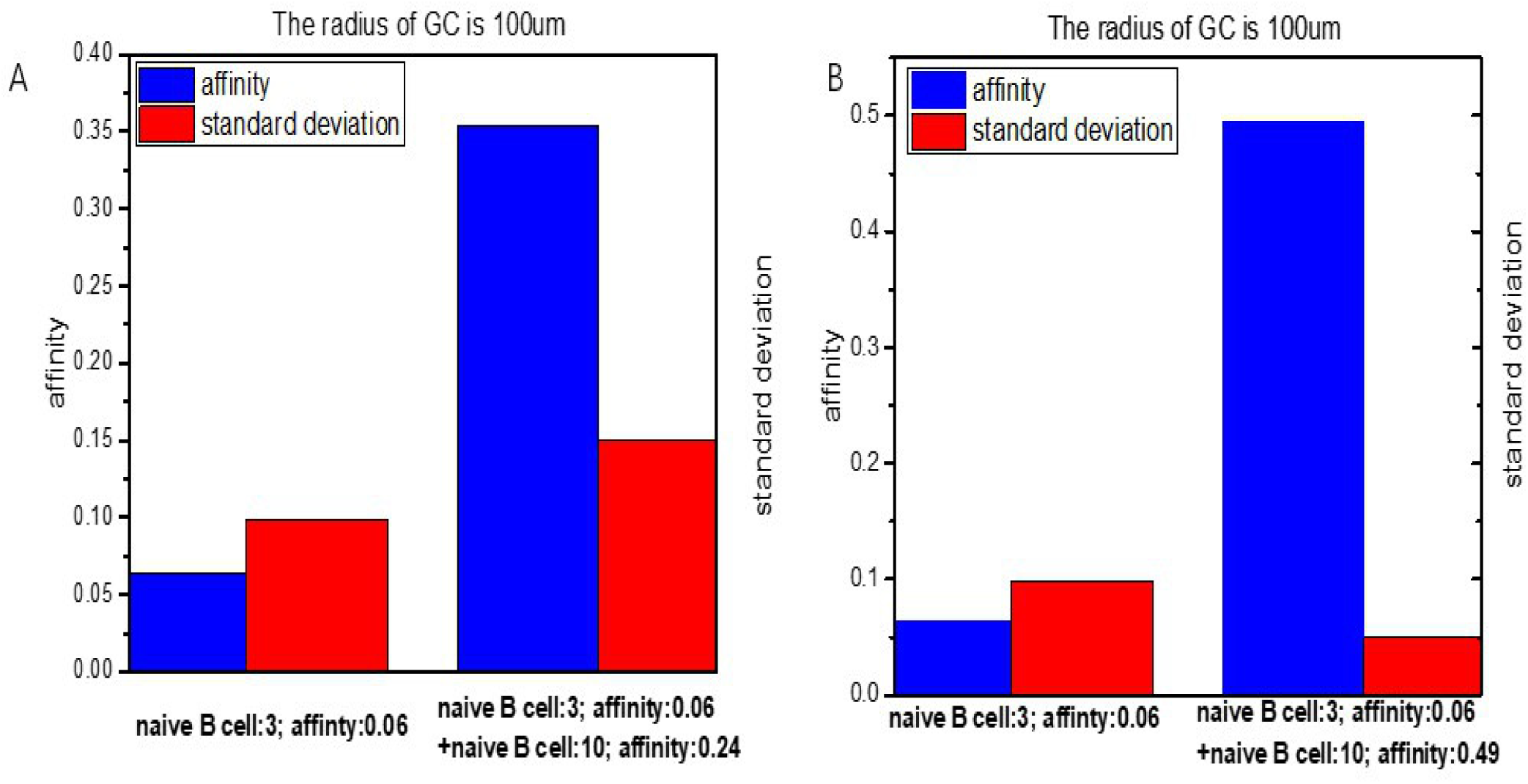
The mean and standard deviation of the affinity of the output B cells. (A): At the initial time, we add 10 B cells with affinity 0.24 in addition to the original 3 B cells with affinity 0.06. (B): Initially, we add 10 B cells with affinity 0.49 instead.

### Simulation on the 3D lattice

All previous simulations are carried out on a two-dimensional lattice. However, realistic biological experiments are performed and observed in three-dimensional space. Therefore, whether the difference in dimensions would significantly change the results, remains to be answered. First, we check the change of reaction rates going from 2D to 3D. In Meyer-Hermann’s article [23], the values of the parameters in the 3D model are directly computed from the 2D. The proliferation rate, differentiation rate, etc. must be multiplied by 3/2 to ensure the results from both simulations are comparable, while the concentrations of FDCs and T cells in GC should be kept constant. As a result, much more FDCs and T cells (= 2700) are needed in three dimensions. However, the scaling of the number of B cells is not so obvious and it is hard to make a direct comparison. So a normalization of the B cell number is needed. As can be seen from Fig.9B, under the same conditions, after the normalization that the number of B cells at each time should be divided by the maximum number of B cells in the whole GC response, the B cells in the 3D simulation seem to be greatly reduced in the later stage of the GC development. It can be seen that the normalization of B cells in the GC in the 3D simulation is nearly 20% less than that in the 2D simulation under comparable conditions. The main reason is that, in our simulation the number of B cells in the 3D is nearly 15 times of that in 2D. As a result even if the gene space and the mutation rate is the same, the mutation range in the gene space of B cells expands. Many B cells stride randomly to a region with very low affinity, so the affinity distribution of B cells changes. As shown in Fig.9D, the proportion of B cells with affinity below 0.05 in 3D is significantly greater than that in 2D. while 0.05 and 0.15, the portion in 2D is greater. As a result, many B cells die quickly because their affinity is too low to get help from T cells. However, if we continue to increase the number of T cells outside the GC from 2700 to 6000 in 3D, arriving at a higher concentration than in 2D, B cells start to get more help from T cells, resulting in a similar result to that of the 2D case. This also explains why large volume of the GC in Fig.4A and Fig.4C leads to a decrease in the affinity.

**Fig 9.**
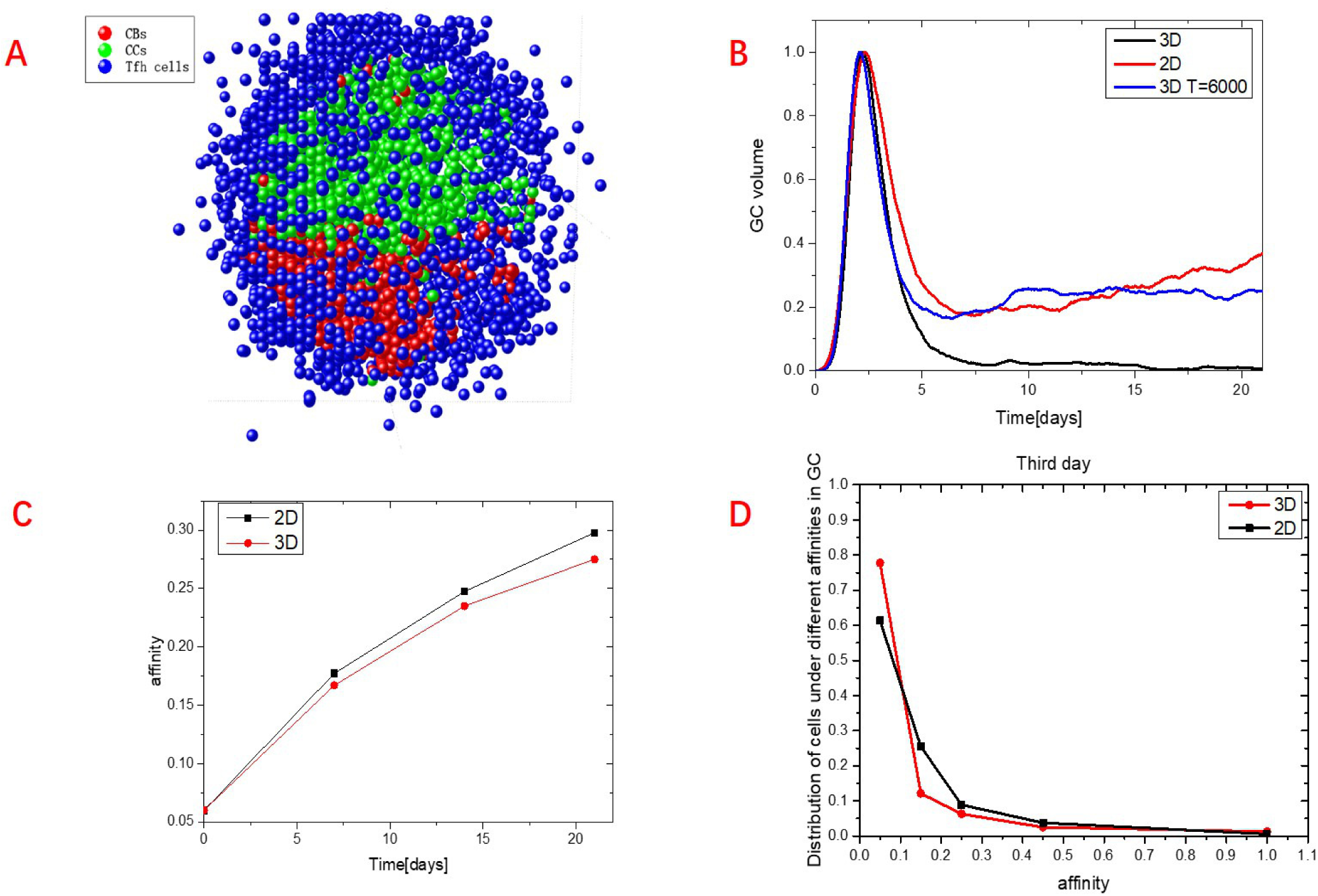
Comparison of the 2D and 3D simulation. (A) a schematic picture of cell distribution in the GC. (B) Time course of the GC volume. In order to compare 3D and 2D, we normalized the data. (C) The average affinity increases with time. (D) Affinity distribution of cells in GC on the third day.

In the 3D simulation, we also check how the radius of the GC alter the affinity maturation. As can be seen from Fig.4B, a phase transition occurs at a radius of 50µm, where the fluctuation rises significantly compared to the 2D case. With a radius of 35µm, the peak number of B cells in the GC is also about 110, being consistent with the results from the 2D simulation (Fig.6D) and the deterministic model. Therefore, if the number of the initial B cells is too small, the resulted few mutations are unable to support a sufficient exploration of the gene space so that affinity maturation could not be achieved before the decay of the GC.

## Discussion

Based on the latest experimental observations, we established a stochastic spatiotemporal model to explore the problem of affinity maturation in adaptive immunity. FDCs present antigens in the form of immune complexes to the naive B cells and allure CCs to compete for antigens on the FDC. The morphology of LZ and DZ plays a very important role in this process. When FDCs are in an unpolarized state, the GC will decay and disappear quickly, and the affinity level of the plasma and memory cells will be very low. By slowly increasing the polarization of FDCs, the structure of the light and the dark zone gradually appears, and the function of affinity maturation gradually recovers, which does show the importance of polarization in the follicle. However, in the process of simulation, because of cell crowdedness, we assume that cellular movement may sometimes be blocked. How much this simulation is consistent with experimental observations remains to be explored.

Next, we explore how different volumes of the affect affinity maturation and whether there is an optimal GC volume. By analyzing the average affinity of differentiated plasma and memory cells in both stochastic and deterministic simulations, we find that with the increase of volume, the average affinity of output cells undergoes a phase transition beyond which a volume may exists that is most favorable for affinity maturation. It turns out that the distance *δ* (*ϕ, ϕ∗*) in the gene space between the activated B-cell and the target epitope of antigen determines the phase transition point and thus the possible optimal volume of the GC. Conversely, different volumes observed in the lymph node may be used as an indicator of this distance if the communication of B cells between GCs could be excluded. Overall, if the number of cell divisions during high frequency mutation is too small to produce enough strains with different affinities, a devastating impact on the subsequent affinity maturation process will be resulted.

The affinity of the naive B cells is very low, being set at 0.06. By the end of the day, the mean affinity of all generated plasma and memory cells rises to a value as high as 0.4 in the current model. However, biological experiments observed that the affinity of plasma cells is about 10 times that of the naive B cells, which suggests other mechanisms at work in the GC, yet to be explored in the future.

## Acknowledgments

This work was supported by the National Natural Science Foundation of China under Grants No. 11775035, and also by the Fundamental Research Funds for the Central Universities with contract number 2019XD-A10.

## A Appendix

### The Basic Stochastic Model

#### A.1 motility and competition

The current model suggests that the movement of the B and T cells can be described as random walks. A cell moves in all directions to the nearest lattice point with the same probability. In a two-dimensional plane, there are four directions to go. However, B cells cannot move to a lattice point occupied by another B cell. When FDCs are polarized in the follicle, the CCs will sense the concentration of CXCL13 secreted by FDCs. The CCs directed movement resulting from the chemoattractant allows the cells leaving the DZ before they die by apoptosis.

To favor the CCs with high affinity, we assume the following mechanism for competition. If one cell is in contact with the a FDC site, and its neighboring cell is not, these two B cells have chance to exchange their positions with probability proportional to the following quantity.

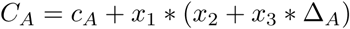

where Δ_*A*_ is the ratio of their BCR affinities. The constant *c*_*A*_ = 2.0 gives the cell’s basic competitiveness, and *x*_1_ = 40.0, *x*_2_ = *−*4*/*3, *x*_3_ = 4*/*3.

In this stochastic model, B cells have the opportunity to contact with T cells by presenting the pMHC molecule to the TCR. One B cell in contact with a Tfh cell may be replaced by a bystander B cell with more pMHC molecules on its surface. The exchange probability is :

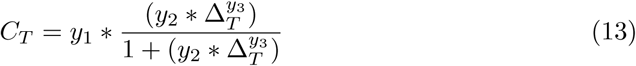

where, Δ_*T*_ is the concentration difference of pMHC and *y*_1_ = 5.0, *y*_2_ = 6.0, *y*_3_ = 0.001.

#### A.2 Clonal proliferation of B cells

In the early stage of the immune response, B cells proliferate rapidly in the DZ, along with SHM. B cells with mutation have different BCR phenotype. The proliferation rate is approximately (9*h*)*−*1 and is controlled by SIP, expressed as:

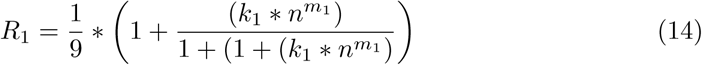

where *n* is concentration of SIP in the B cells, *m*_1_ = 6, *k*_1_ = 4.3519 × 10*−*4. At the initial stage, since the seeder B cells contain a high concentration of SIP (*cl* = 1000), they rapidly divide. SIP and other proteins are disseminated randomly to daughter cells after B cell proliferation. Over time, the cell division rate will decrease with the decrease of intracellular SIP concentration, so B cells need help signals from Tfh cells to synthesize more SIP.

In the simulation, the rate of B cell divisions depends not only on *R*_1_, but also on the number of vacancies in a neighborhood of the cell. Because each grid point can only be occupied by a cell of the same type, the number of B cells may seriously affect the rate of its division. For example, when there are eight B cells around a B cell, this B cell will not be able to divide if we assume that each one is fixed to the occupied lattice site. But this assumption is apparently not consistent with experimental observation. In fact, B cells can push other cells away for division. In order to cope with this problem, we assume that in a neighborhood with a radius of 100 *µ* m, if there are vacancies, the B cell may divide normally and the newly generated daughter-cell will be randomly put into to a vacancy in this neighborhood.

#### A.3 B cell differentiation

There is still no clear conclusion on how CBs differentiate into CCs. It has been stated based on experimental observation that the transition is independent of location. It might be triggered by LZ-derived signals such as the receipt of the T cell signal. In our model, the differentiation rate is controlled by the concentration of SIP. When the concentration of SIP in CBs was low, CBs began to differentiate into CCs. CCs, neither proliferating nor mutating express BCR on the surface of the cell membrane. They are in an activated state of apoptosis and have a life span of about 6 hours. In the model, the concentration of SIP in CBs determines the differentiation rate.

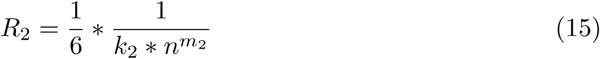

where *n* is the concentration of SIP in the B cells, *m*_2_ = 5 is Hill’s coefficient, *k*_2_ = 0.0102. Proliferation of B cells leads to a decrease in the SIP concentration, which not only promotes the differentiate of CBs into CCs, but also increases the sensitivity of B cells to apoptosis. Therefore, the CCs with higher affinity are able to enter LZ to capture antigens and present them to T cells to reduce the rate of apoptosis. After the CCs receive the help signal from the Tfh cells, they start to synthesize and accumulate the SIP. These selected CCs may differentiate into CBs and return to the dark zone, where the mutation and selection is restarted. The differentiation rate *R*_3_ of CCs to CBs is :

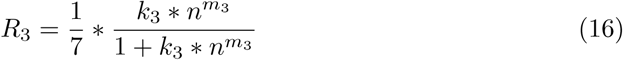

where *n* is the concentration of the SIP, *m*_3_ = 4 is the Hill’s coefficient, *k*_3_ = 0.0011.

#### A.4 Apoptosis of cells

According to experiments in vivo, CCs can survive in the LZ for about 6 *∼* 16 hours. Because SHM makes BCR affinity differ from one another, BCR goes through the selection of antigens presenting on FDCs and only those B cells expressing high affinity antibodies can effectively seize antigens from FDCs. Those with low affinity will be eliminated due to their high sensitivity to apoptosis. It is worth noting that the B cell apoptotic rate is essential in the regulation of GC during the immune response. If the rate of apoptosis of B cells is too high, B cells will all die in a short time. A high immunity level is not achievable. Conversely, if the rate of apoptosis of B cells is too slow, B cells will occupy most lattice points in the GC. As a result, the SHM cannot proceed smoothly, and finally slow down of BCR affinity maturation. During the simulation, the state vector of apoptotic B cells will be directly deleted and removed from the computer memory. In the model, we assume that the apoptotic rate increases with the decrease of SIP, so the rate *R*_4_ is :

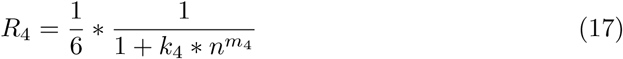

where *n* is the concentration of the SIP, *m*_4_ = 5 is the Hill’s coefficient, *k*_4_ = 0.05.

#### A.5 Interaction between CCs and FDCs

Follicular dendritic cells (FDCs), the most powerful antigen-presenting cells (APCs) in GC, efficiently absorb, process, and present antigens. The presented antigen binds to BCR to form an immunological synapse and plays an important role in activating B cells. B cells with high affinity will endocytosis antigens, and then within the B cell, antigens will be decomposed into polypeptide fragments, part of which then bind to the MHC class II molecule to form a complex and are transferred to the B cell surface. These complexes can be recognized by Tfh cells. In the model of this paper, CCs enter the lZ of GC, and may undergo three states: “not selected”, “contact with antigen” and “antigen selection”:

##### A.5.1 Unselected

CCs that are not in contact with antigens on FDCs will not be recognized by Tfh cells. Therefore, the concentration of SIP in B cells do not increase. The apoptosis was highly sensitive to lower concentration of SIP in B cells, so CCs will be eliminated by apoptosis at a rate of *R*_4_.

##### A.5.2 Contact with antigens

The “contact with antigens” CCs indicate that BCR has successfully contacted with antigens in FDCs, but there is no endocytosis antibody – antigen immune complex. We assume that the binding rate is related to the affinity of the BCR, and the CCs remain bound to the antigens for about 5min. The antigen binding rate *R*_5_ is recorded as:

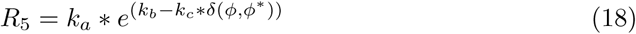

where *k*_*a*_ = 2.696, *k*_*b*_ = 0.8, *k*_*c*_ = 0.25. *δ* (*ϕ, ϕ∗*) represent the distance between the BCR and the antigen in the gene space 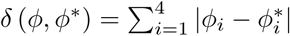.

##### A.5.3 Selected by antigens

CCs selected by antigens indicate that BCR successfully ingested antigens from FDCs. In this model, the reaction rate of B cells to take up antigens is related to the affinity of BCR. It is given as:

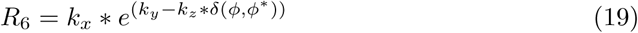

where *k*_*x*_ = 0.0056, *k*_*y*_ = 5.6, *k*_*z*_ = 0.7. With the highest affinity, a CC will take up one Ag about every 4min. As you can see from Fig.2A, the rate of uptaking Ag from FDCs is higher when the affinity is higher.

#### A.6 Interaction between CCs and Tfh cells

Professor Hai Qi used TIRF microscopy to observe Tfh cells entangled with B cells. During T-B entanglement, two co-stimulatory molecules of ICOS-ICOSL and CD40L-CD40 constitute a positive feedback network for improving affinity maturation of BCR. Recently, they found the role of another pair PD-1-PD-L1, the programmed cell death-1(PD-1) expressed by Tfh cells that controls the location and function of Tfh cells. It may dampen TCR signaling and thereby reduce the ligand sensitivity of Tfh cells, which should increase the overall stringency of selection in GCs. In real biological experiments, PD-L1 deficient B cells can get more help from Tfh cells. Based on these data, it was shown that the interaction of PD-1 and PD-L1 could also increase the affinity of BCR. These findings should be reflected in the stochastic model. Similarly, during the simulation, CCs after uptaking antigens could dwell in three different states: “not selected by T cells”, “contacted by T cells” and “selected by T cells”:

##### A.6.1 Not selected by T cells

CCs that are “not selected by T cells” indicate those that have not obtained help signal from T cells since entering the LZ. Therefore, these CCs will be difficult to survive in the LZ unless they contact with a T cell during the wandering in the LZ.

##### A.6.2 Contact with T cell

Once CCs have access to a T cell site, they will have chance switching to the state of contact with T cell. However, in this situation, no help signals are transmitted from T cells to B cells. The binding rate is regulated by the concentration of antigen taken by CCs and the concentration of PD-L1 on the surface of the cell. If it has too much PD-L1, the probability of binding will be close to zero. The T-B binding rate *R*_7_ is given as:

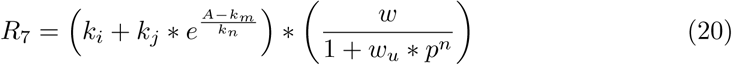

where A is the concentration of antigens, *k*_*i*_ = 10, *k*_*j*_ = 2, *k*_*m*_ = 10, *k*_*n*_ = 30. p is the concentration of PD-L1,*w*_*u*_ = 0.4, *n* = 5, *w* = 10. The binding sites for the antigen and the PD-L1 are independent of each other, so the total rate *R*_7_ is a product of these two independent factors.

##### A.6.3 Selected by Tfh

CCs selected by T cells are mainly those with a high concentration of peptides on the cell surface. When selected by T cells, the MHC complex and ICOS-ICOSL work coherently to make CCs obtain help signals from the T cells. In the model constructed in this paper, the rate of B cells to get the help signal *R*_8_ is

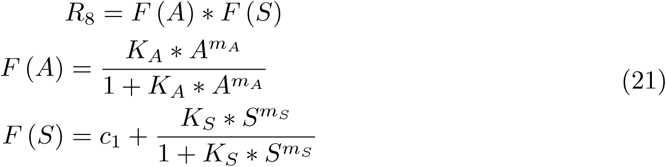

where A and S are the concentration of antigens and ICOSL. The two factors *F* (*A*) and *F* (*S*) indicates the action from the antigen and ICOSL, respectively. They are assumed to work independently here. *m*_*A*_ = 8 and *m*_*S*_ = 2 are the Hill’s coefficient. Constant: *K*_*A*_ = 6 × 10^*−*4^, *K*_*s*_ = 4, *c*_1_ = 18.5, *c*_2_ = 4. The T-B interaction allows CCs procuring help signals from T cells, and then CCs can synthesize SIP with a synthesis rate of 1. Therefore, CCs with higher affinity have lower apoptotic rate, and thus those B cells survive better in the GC. The number of help signals will gradually decrease.

#### A.7 Cell output

CCs with high affinity in LZ is more likely to undergo multiple selection and mutation cycles, which may differentiate into plasma or memory cells, and leave GC within 7 hours. In the model, the rate R9 at which CC differentiates into plasma or memory cells is determined by the concentration of SIP.

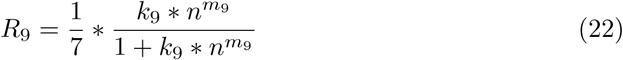

where n is the concentration of the SIP, *m*_9_ = 4 is the Hill’s coefficient and *k*_9_ = 0.0011. CCs can only differentiate into plasma or memory cells when the concentration of SIP is high enough in the cell. At the same time, according to the model, the probability of “CC differentiation into CB” is four times that of “CC differentiation into plasma or memory cell”. Plasma or memory cells will directly leave GC. The antibody secreted by plasma cells is assumed not affecting the immune response in GC, and the corresponding cell state vector will be deleted directly.

